# Developmental tuning of mineralization drives morphological diversity of gill cover bones in sculpins and their relatives

**DOI:** 10.1101/437749

**Authors:** Eli G. Cytrynbaum, Clayton M. Small, Ronald Y. Kwon, Boaz Hung, Danny Kent, Yi-lin Yan, Matthew L. Knope, Ruth A. Bremiller, Thomas Desvignes, Charles B. Kimmel

## Abstract

The role of osteoblast placement in skeletal morphological variation is relatively well understood, but alternative developmental mechanisms affecting bone shape remain largely unknown. Specifically, very little attention has been paid to variation in later mineralization stages of intramembranous ossification as a driver of morphological diversity. We discover the occurrence of specific, sometimes large regions of nonmineralized osteoid within bones that also contain mineralized tissue. We show through a variety of histological, molecular, and tomographic tests that this “extended” osteoid material is most likely nonmineralized bone matrix. This tissue type is a significant determinant of gill cover bone shape in the teleostean suborder Cottoidei. We demonstrate repeated evolution of extended osteoid in Cottoidei through ancestral state reconstruction and test for an association between its presence and habitat differences among species. Through measurement of extended osteoid at various stages of gill cover development in species across the phylogeny, we gain insight into possible evolutionary developmental origins of the trait. We conclude that this finetuned developmental regulation of bone matrix mineralization reflects heterochrony at multiple biological levels and is a novel mechanism for the evolution of diversity in skeletal morphology. This research lays the groundwork for a new model in which to study bone mineralization and evolutionary developmental processes, particularly as they may relate to adaptation during a prominent evolutionary radiation of fishes.

## Introduction

Understanding the genetic, developmental, and evolutionary mechanisms that underlie morphological variation is an overarching aim in many branches of biology. Bone shape and size constitute a fundamental component of morphological diversity, and a rich body of work using vertebrate animal models - primarily mouse and zebrafish - has identified developmental pathways that, when disrupted, significantly alter skeletal structure (see Clouthier and Schilling (2004) for a review of major craniofacial effectors). Large-effect perturbations introduced in these studies affect processes such as pharyngeal arch patterning through Endothelin-1-dependent signalling cascades (Kurihara et al. 1994; Neuhauss et al. 1996; Ruest et al. 2003), and Krox-20-dependent skeleton-wide ossification (Levi et al. 1996), so they often have severe fitness consequences and are unlikely to explain most of the variation we observe in nature. An important goal, therefore, is to identify developmental mechanisms that permit specific and less disruptive (but still morphologically significant) changes to bone shape relevant at the population and species levels. Such variants likely manifest during an extended period of development, resulting in a reshaping or resizing of bones we observe and recognize as homologous in different species. Our previous work in zebrafish, for example, revealed that modular regulation of *osteoblast placement* within a skeletogenic mesenchymal condensation is a critical determinant of bone shape (Kimmel et al. 2010). Another osteoblast-specific mechanism of shape modification occurs by increasing the *number of osteoblasts* in a region on the edge of the bone, which can be accomplished by cell recruitment through fate switching (Takada et al. 2009), by cell migration (Fukuyama et al. 2004), or by local osteoblast or pre-osteoblast proliferation (Huycke et al. 2012; Kim et al. 2012).

While the spatiotemporal distribution of osteoblasts is clearly an important determinant of bone shape, modification of later events during osteogenesis may require fewer perturbations to core regulatory networks important during skeletal development. A more modular “fine tuning” of late stages in bone development, therefore, might be especially accessible to evolutionary processes like natural selection (Wagner 1996; Brakefield 2006; Mengistu et al. 2016). During intramembranous ossification, demarcation of the bone tissue precursor is based on the spatial distribution of osteoblasts, but secretion of a collagen-proteoglycan osteoid matrix must follow (Franz-Odendaal et al. 2006). This osteoid tissue is transient, and provides the organic matrix for mineralized bone, in part through the expression of extracellular matrix proteins like Fibronectin, Type I Collagen, Bone Sialoprotein, Osteopontin, and Osteocalcin (Ducy et al. 1997; Cowles et al. 1998). It is entirely conceivable that a bone’s effective shape, and ultimately its function, could be determined by the extent to which mineralization of the osteoid matrix occurs, serving as a potentially important mechanism for the developmental tuning of skeletal shape and ultimately the generation of variation at a macroevolutionary scale.

We have observed widespread variation across acanthomorph fishes in the shape of the mineralized opercle (OP), the primary gill cover supporting bone (Kimmel et al. 2017). Based on our study of OP morphospace, some of the most striking variation over a relatively short phylogenetic distance occurs in fishes of the suborder Cottoidei, which includes sculpins, sandfishes and snailfishes (Smith and Busby 2014). In this article we refer to all fish in the suborder Cottoidei as “cottoids.” To avoid confusion, when referring to the cottoid superfamily Cottoidea (Jordaniidae, Rhamphocottidae, Scorpaenichthyidae, Agonidae, Cottidae, and Psychrolutidae), we will use the full superfamily name. The Cottoidea, which include “sculpins” and “poachers,” underwent a morphologically diverse radiation of species as recently as the Miocene (David 1943) and have evolved to occupy a range of environments, including multiple transitions from subtidal marine to intertidal (Knope and Scales 2013) and freshwater (Knope 2013; Goto et al. 2014) habitats. Indeed, their rate of diversification has been calculated to be on par with that of cichlids (Near et al. 2013), a canonical group for the study of rapid diversification and speciation (Salzburger et al. 2005). We find that sculpin lineages differ from one another dramatically with respect to the presence of mineralized material in the central, distal region of the OP, and to a much lesser extent in regions of neighboring bones including the subopercle (SOP) and the interopercle (IOP). This variation is likely of functional relevance given the important roles of gill cover bones in respiration (Hughes 1960), mouth opening, and predator defense (Anker 1974), and given associations between variation in OP shape and ecological variables in Lake Tanganyikan cichlids (Wilson et al. 2015), ariid catfishes (Stange et al. 2016), threespine stickleback (Kimmel et al. 2008; Kimmel et al. 2012), and icefishes (Wilson et al. 2013).

Here we study this exceptional diversity in mineralized shape of the OP bone among cottoid fishes (and several outgroups) and describe a prominent feature we call “extended osteoid,” bone matrix that makes up large regions of certain bones and is dynamic, pliable, and calcium-deficient in nature. We evaluate the developmental and evolutionary significance of extended osteoid by addressing several important questions about its properties and diversity: 1. Are the large, membranous regions of gill cover bones observed across species biologically classifiable as osteoid? 2. Is the majority of OP mineralized shape variation among species explained by extended osteoid? 3. Have largely nonmineralized OPs evolved multiple times across the cottoid phylogeny, and beyond? And 4. How does the developmental timing of the appearance of extended osteoid vary among species? In answering these questions we propose the phenomenon of extended osteoid as a mechanism for explaining the major OP diversity of the Cottoidea and present a new vertebrate model in which to study the role of bone mineralization in development and evolution.

## Materials and Methods

### Collection and identification of specimens

The majority of individuals represented in this study were obtained as loans from other researchers, scientific collections, or aquaria. These include the Oregon State Ichthyology and Burke Museum collections, the Vancouver Aquarium, the Oregon Coast Aquarium, bycatch from other researchers at the University of Oregon, and fish from previous studies (Knope 2013; Knope and Scales 2013; Kimmel et al. 2017). All species considered in our current study, the sources of the material, and the analyses based on them, are included in Supplementary File 1. Our survey included seven families from the suborder Cottoidei (Liparidae, Jordaniidae, Rhamphocottidae, Scorpaenichthyidae, Agonidae, Cottidae, and Psychrolutidae), plus six outgroup perciform families (Hexagrammidae, Zoarcidae, Pholidae, Gasterosteidae, Sebastidae, and Centrachidae).

We also collected some of our own specimens through hand netting and trapping in Oregon streams and tidepools. All fish taken live by the authors were captured, euthanized using a lethal dose of MS-222 (tricaine), and preserved in ethanol or 4% paraformaldehyde (in situ hybridization samples only). This study was approved by the University of Oregon IACUC (Protocols #17-28 and #10-26), and all live animals used for this study were treated in accordance with this IACUC protocol.

Fish were morphologically keyed using Miller and Lea (1972) in addition to Markle (1996). All initially ambiguous species designations were confirmed by BLAST searches based on cytochrome b mitochondrial DNA and s7 intron sequences, commonly used markers for understanding sculpin species relationships (Grachev et al. 1992; Ramon and Knope 2008). We extracted DNA from fin clips stored in 95% ethanol or stored at -80 °C using a preciptation-free, lysis-only DNA extraction protocol (Westerfield 2000). Samples were PCR-amplified for mitochondrial cytochrome b (GLUDG-L: 5’-TGACTTGAARAACAYCGTTG-3’ and CB3-H: 5’-GGCAAATAGGAARTATCATTC-3’ for marine species and L14724: 5’-GTGACTTGAAAAACCACCGTT-3’ and H15915: 5’-CAACGATCCGGTTTACAAG-3’ for Cottus) and the first intron of the nuclear S7 ribosomal protein (S72F: 5’-TCTCAAGGCTCGGATACGTT-3’ and S74R: 5’-TACTGAACATGGCCGTTGTG-3’ for all fish), followed by clean-up and Sanger sequencing.

### Sample preparation for morphological assay

#### Staining

Fish used for morphological analysis were stored in 70% or 95% ethanol until they were stained using the Alcian Blue and Alizarin Red double stain protocol (Walker and Kimmel 2007). Alcian Blue stains cartilage and other tissues via binding to polysaccarides, and Alizarin Red stains mineralized bone tissue by binding directly with calcium-containing compounds. The stained bones were then stored in 50% glycerol with 0.1% potassium hydroxide and a small amount of thymol.

#### Dissection

Using forceps, the dentaries were disarticulated along with the premaxillae and ceratohyals. The hyomandibula was disarticulated from the cranium, freeing the facial bones. The bones were then completely disarticulated and cleaned. This method was found to be most consistent while preserving all hard and soft tissues of bone elements. Very seldom, trypsin was used to help soften the tissue.

#### Imaging

Dissected bones were flat-mounted and imaged using bright-field, Nomarski differential interference contrast (DIC) and incident fluorescent lighting. Using Adobe Photoshop we simplified the bones to detailed silhouettes, both including and excluding the extended osteoid portions. These silhouettes formed the basis of morphometric (shape) analyses and “proportion osteoid” calculations (see below), which were carried out using pixel counts.

### Compositional analysis of extended osteoid

#### In situ hybridization

We designed oligonucleotide probes for the transcription factor *sp7* (also known as *osterix*) using a conserved region in an alignment of coding sequences (Supplementary File 2) from the assembled whole-body transcriptomes of *Cottus perplexus*, *Oligocottus maculosus*, and *Clinocottus globiceps*. These transcriptome assemblies were generated with 100 nt paired-end Illumina reads using Trinity (Grabherr et al. 2011), and they are part of a separate, forthcoming study. Probe design followed the methods of Albertson et al. (2010). In zebrafish, *sp7* is expressed robustly in osteoblasts and shows stronger specificity than other osteoblast-expressed genes such as *runx2a* and *runx2b* (Li et al. 2009), justifying its use as a reliable marker for bone tissue from the onset of mineralization.

Upon euthanasia, heads of subadult *O. maculosus* and *C. globiceps* specimens were fixed in 4% PFA and incubated for ~24 hours at 4° C, after which they were washed twice in PBT (1xPBS, 0.2% Tween-20). Each head was then soaked in one milliliter for three minutes each in a progression of 25% methanol : 75% PBT; 50% methanol : 50% PBT; 75% methanol : 25% PBT before soaking twice in 100% methanol. Heads were stored at -20o C in 100% methanol. We then dissected out the portion of the operculum containing the OP and SOP and performed *in situ* hybridization as described in (Yan et al. 2005).

#### Histology

We fixed heads of juvenile Cottus perplexus, and two eelpouts (*Ophthalmolycus amberensis*, and *Lycenchelys tristichodon*) in 4% PFA, decalcified, embedded in paraffin, and cut 10 μm cross sections. We stained the slide-mounted sections using the sensitive tetrachrome procedure described by Ralis and Watkins (1992), which differentiates between osteoid and mineralized bone.

#### Micro CT

MicroCT scanning was performed using a vivaCT40 (Scanco Medical, Switzerland). The OP from *Scorpaenichthys marmoratus* was scanned using the following settings: 38 μm isotropic voxel size, 55 kVp, 145 mA, 1024 samples, 500 proj/180 °, 200 ms integration time. High-resolution scans of the OP from *Oligocottus maculosus* were acquired using the following settings: 10.5 μm isotropic voxel size, 55 kVp, 145mA, 2048 samples, 1000 proj/180 °, 200 ms integration time. DICOM files of individual fish were generated using Scanco software, and analyzed using FIJI.

### Morphological and evolutionary analyses

#### OP Shape analysis

We used the extended eigenshape method of shape analysis, developed by MacLeod (1999); (MacLeod 2012), who kindly contributed software in the form of unpublished Mathematica notebooks that included all of the required procedures. The analysis is two-dimensional, appropriate for the flattened form of the OP. We initially examined a set of 122 samples spanning 13 perciform families, which included some species replicates and juvenile and subadult stages. For the current study, however, we culled this set to include only adults, and in the interest of equal taxon sampling, only a single individual per species. Using this culled set of 43 species, we placed landmarks at three specific regions of the bone edge: 1. at the middle of the joint socket for the articulation the OP makes with the hyomandibula, 2. at the tip of the ventral spur, and 3. at the tip of the posterior spur. These landmarks separated three segments along the bone outline that we demarcated by semi-landmarks, placed at equal intervals to one another along the mineralized portion of the bone edge, to examine shape variation within these segments: 1. 11 semi-landmarks along the ‘anterior segment’ between landmarks 1 and 2; 2. 57 semi-landmarks along the ‘ventral-posterior segment’ between landmarks 2 and 3, and 3. 19 sem-ilandmarks along the ‘dorsal-posterior segment’ between landmarks 3 and 1. These xy coordinates (90 in all) captured >90% of shape variation of the OP perimeter. The coordinates were Procrustes-transformed to remove variation due to size and orientation, and the resulting shape coordinates subjected to principal component analysis (PCA) to identify and characterize the principal shape changes contributing to morphological diversity. All downstream analyses were conducted using version 3.3.2 of the R statistical language (Team 2016). To assess the relationship between proportion of extended osteoid and the first principal component scores we performed standardized major axis regression using the R package SMATR (Warton et al. 2011). This regression analysis was performed using phylogenetically independent contrasts (Felsenstein 1985) calculated for both variables using the pic function from the R package APE (Paradis et al. 2004).

#### Ancestral state reconstruction (ASR) and trait-habitat association analysis

We performed binary (“fork” or “fan”) ASR for OP shape among adult fish from the 43 species featured in the extended eigenshape analysis (see above), using maximum likelihood and parsimony approaches. The topology we used was primarily based on Cottoid relationships from Smith and Busby (2014), but also Baumsteiger et al. (2012), Goto et al. (2014), Buser and Andres Lopez (2015), and outgroup relationships from Near et al. (2013). In the absence of a chronogram, we calculated branch lengths according to Grafen’s method (Grafen 1989). We used the ace function from the R package APE (Paradis et al. 2004) to infer discrete ancestral states using an all-rates-equal (ER) model to calculate marginal likelihoods. Parsimony-based inference was carried out using the MPR function of APE, in accordance with the approach of Narushima and Hanazawa (1997). Using all of the PCA species except two for which we could obtain a reliable estimate of the proportion of nonmineralized OP bone (*Clinocottus globiceps* and *Enophrys bison*), we performed continuous-character ASR. This was carried out again using the ace function of APE, assuming a Brownian motion model of character evolution.

We also tested whether habitat is associated with the proportion of extended osteoid across all lineages of the tree (excluding the derived freshwater outgroup Centrachidae), using generalized least squares (gls) taking into account phylogenetic correlation. We estimated a phylogenetic correlation matrix under Brownian motion using the *corBrownian* function of APE, which we then used to fit a gls linear model describing the relationship between trait values and environment. The gls model was fit using the function gls from the R package nlme (Pinheiro 2017). Similarly, to test whether the binary trait of “fan/fork” OP morphology was associated with habitat we carried out phylogenetic logistic regression using the *phyloglm* function from the R package phylolm (Ho and Ane 2014). Habitat designations for species, both tested as a three-level factor (deep marine/shallow marine/freshwater or estuary) and a two-level factor (deep/shallow), were based on data from fishbase.org and Knope and Scales (2013). Intertidal and “transitional” marine habitats and freshwater habitats were considered “shallow,” and subtidal marine habitats were considered “deep.”

## Results

### Extended osteoid is a prominent feature of cottoid opercular bones

#### Extended osteoid is common in cottoid fishes

18 of the 19 genera we surveyed within the Cottoidei exhibited at least some flexible, membranous tissue within the three primary bones that make up the operculum, or “gill cover.” This membrane appeared to lack calcified matrix, as indicated by clear regions negative for Alizarin Red staining (Fig 1A-B; Fig S1). We named the membrane “extended osteoid,” to reflect its likely makeup as a long lasting, dynamically regulated osteoid matrix which appears to undergo a delay in the final stages of intramembranous ossification (Cowles et al. 1998). With the possible exception of Agonidae (poachers), all cottoids we surveyed possessed this extended osteoid as part of the subopercle (SOP), where it lines the ventral and to a lesser extent dorsal edges of the SOP’s posterior projection. Additionally, some cottoid species demonstrated osteoid membrane located at the ventral edge of the interopercle (IOP). We observed a remarkable degree of variation across the superfamily Cottoidea in the amount of extended osteoid present in the opercle (OP), ranging from a complete or nearly complete absence of this tissue type (a fully Alizarin Red-stained “fan” shape) to a “fork” shape in which the entire central portion of the OP was unstained (Fig 1 B and A, respectively). We also surveyed nine (non-cottoid) outgroup genera from the order Perciformes (Betancur-R et al. 2017), representing the families Hexagrammidae, Zoarcidae, Pholidae, Gasterosteidae, Sebastidae, and Centrachidae. Of these outgroups we found evidence for noncalcified, membranous bone only in the OPs of the two zoarcids we examined, the eelpout species *Lycenchelys tristichodon* and *Ophthalmolycus amberensis* (Fig S2A-E).

**Figure 1.**
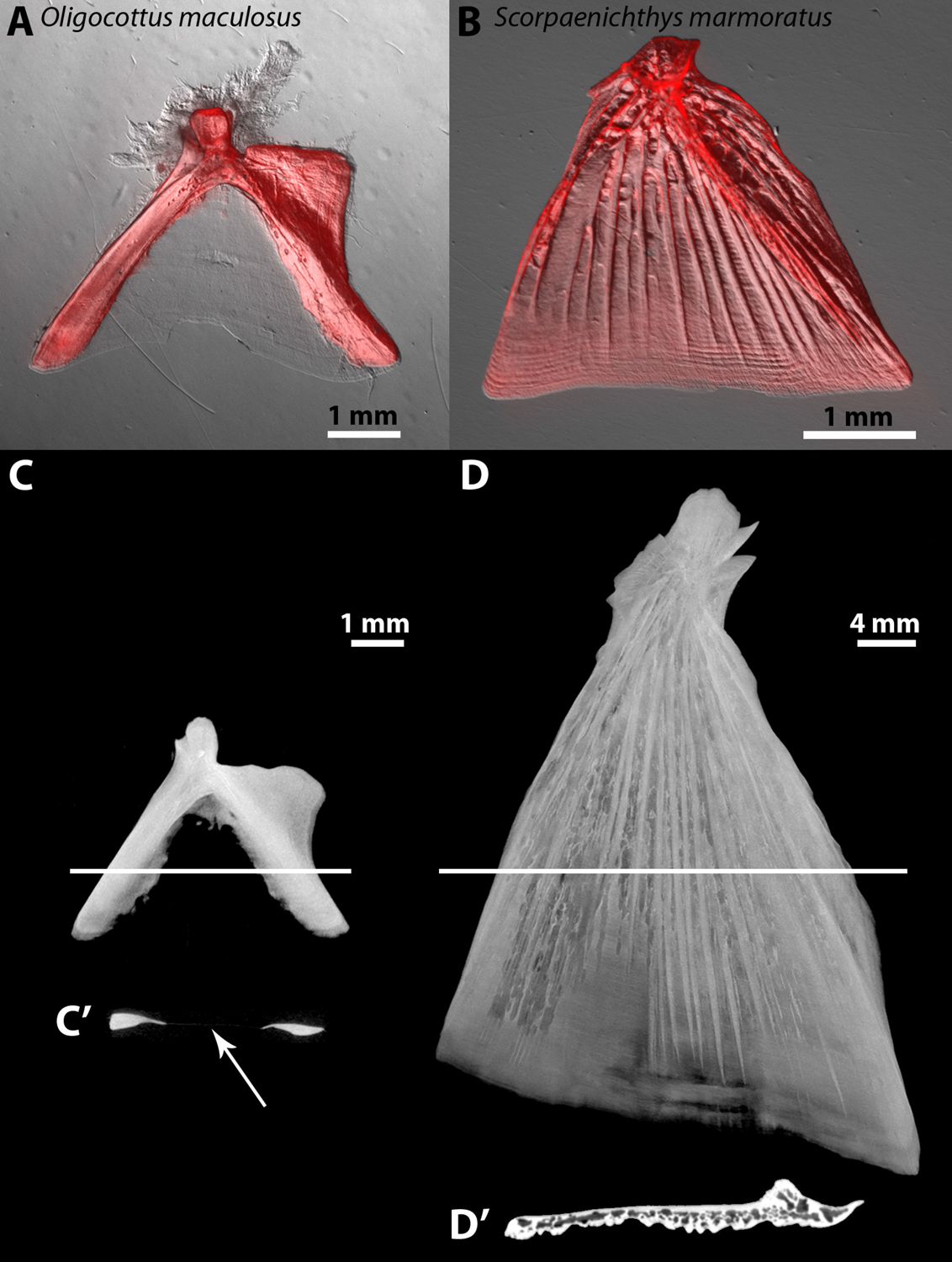
Extended osteoid shows a lack of calcification consistent with known umineralized tissues. *A*. An adult *Oligocottus maculosus* OP stained with Alizarin Red, showing lack of stain in the region of extended osteoid. *B*. A juvenile *Scorpaenichthys marmoratus* OP stained with Alizarin Red, showing full calcification throughout the bone. *C*. An adult *O. maculosus* OP scanned using micro-computerized tomography (μ-ct) clearly shows mineralized and nonmineralized (extended osteoid) regions, and a computed cross section of the scan (*C*’) shows a nearly undetectable tissue layer in the extended osteoid region (arrow). *D*. An adult *S. marmoratus* OP scanned using micro-computerized tomography (μ-ct) clearly shows consistent mineralization throughout, albeit with heterogeneous density typical of reticular bone, as seen in the virtual cross section (*D*’).

#### Extended osteoid is fundamentally distinct from mineralized bone

Alizarin Red staining revealed that whereas calcified portions of bones stained robustly, extended osteoid regions did not take up stain noticeably more than the background levels present in other tissue types known to be nonmineralized, such as connective tissue (Fig 1A). Furthermore, micro-computerized tomography (μ-ct) scanning of an *Oligocottus maculosus* OP, for example, showed the extended osteoid to display extremely low radiopacity (Fig 1C), as confirmed by an image constructed perpendicular to the plane of the bone (Fig 1C’), and suggesting the presence of very thin, likely nonmineralized tissue between the two heavily mineralized “struts.” Conversely, the species *Scorpaenichthys marmoratus* (a more basal lineage in Superfamily Cottoidea) showed little, if any, OP extended osteoid. We observed positive Alizarin Red staining throughout most of its OP, except for a very thin strip of about 30μm at the leading edge (Fig 1B). A (μ-ct) scan for a *S. marmoratus* OP also suggested clear mineralization throughout most of the bone (Fig 1D-D’).

Ralis-Watkins staining, which is a standard method for differentiating mineralized bone from osteoid (Ralis and Watkins 1992), confirmed that in *Cottus gulosus* (Fig 2A-B), and two non-cottoid (eelpout) species (Fig S2C-D’), extended osteoid was not simply extremely thin, canonically mineralized bone. The extended osteoid stained a deep blue (as one expects for osteoid), while the surrounding mineralized bone stained red (as one expects for mineralized bone).

**Figure 2.**
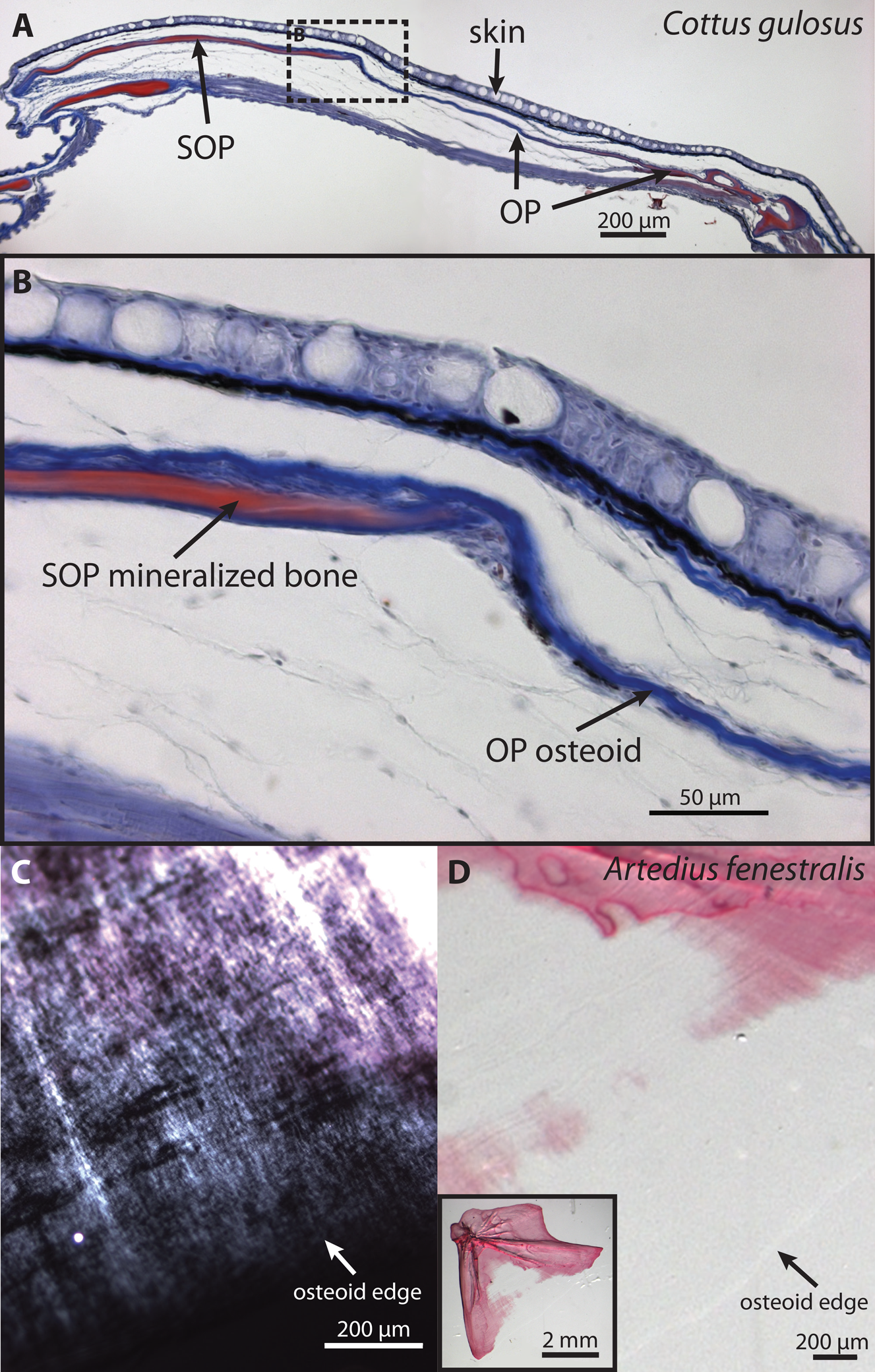
Ralis-Watkins staining and “growth bands” confirm the osteoid nature of the nonmineralized membrane in OPs of two “fork-bearing” sculpins. *A*. A cross section of a Ralis-Watkins stained *Cottus gulosus* gill cover, with the mineralized portion of the OP and SOP in red and the extended osteoid in blue, showing a lack of mineralization in the extended osteoid region. The two OP arrows straddle the transition between nonmineralized and mineralized regions of the OP. *B*. Higher magnification of the same image, corresponding to the dashed rectangle in *A*. *C*. Alizarin-stained *Artedius fenestralis* OP showing banding continuous between the mineralized (pink) and nonmineralized (gray) portions of the bone, visualized with Nomarski differential interference contrast microscopy. *D*. Same image with lower magnification and cross-polarized light. Inset is a view of the entire bone.

#### Structural and cellular evidence supports extended osteoid as true osteoid

Nomarski and cross polarized light imaging revealed a developmental signature of incremental banding (“growth rings”) in extended osteoid regions of the OP, which was in phase with banding of the adjacent mineralized bone (Fig 2C-D). This pattern is consistent with an ordered structure of pre-mineralized bone matrix (Reznikov et al. 2018) and known circadian “periodicity” in osteoblast proliferation (Fu et al. 2005), and therefore an expectation for osteoid independent of the mineralization process. The banding we observed was synchronized and continuous between mineralized and nonmineralized regions of the OP (Fig 2D), suggesting that extended osteoid initially forms in conjunction with the neighboring osteoid that ultimately follows the more standard mineralization trajectory.

*In situ* hybridization in juvenile *Oligocottus maculosus* and *Clinocottus globiceps* revealed robust transcription of *sp7* at the growing edge of OP extended osteoid (Fig S3). *sp7* is expressed by differentiating osteoblasts with high specificity at the osteogenic front of the OP in zebrafish (Li et al. 2009; Huycke et al. 2012). The lining of extended osteoid with functional osteoblasts suggests that this tissue is indeed a form of osteoid.

### Extended osteoid explains the major axis of OP shape variation in cottoid fishes and beyond

As mentioned, species from the superfamily Cottoidea (Smith and Busby 2014) fell along a continuum from fan-shaped to fork-shaped OPs, when considering the calcified (Alizarin Red-stained) bone outline (Fig S4). We performed a shape outline-based principal components analysis (PCA) including this monophyletic group and seven additional families, using only mineralized OP outlines (excluding extended osteoid) to define “effective” shape. The first principal component (PC1) of mineralized OP PCA explained 56.28% of the total shape variation, and it mostly separated “fan-like” from “fork-like” shapes (Fig 3A). OP outline regions loading most heavily on PC1 were concentrated at the center of the OP ventral-posterior margin (Fig 3B), where most of the variation regarding extended osteoid resided. Outline regions loading most heavily on PC2 and PC3 were concentrated at the joint region and dorsal-posterior edge of the OP (Fig 3B), locations of attachment for the dilator operculi and levator operculi muscles, respectively (Yabe 1985), suggesting important functional shape variation perhaps independent of extended osteoid.

**Figure 3.**
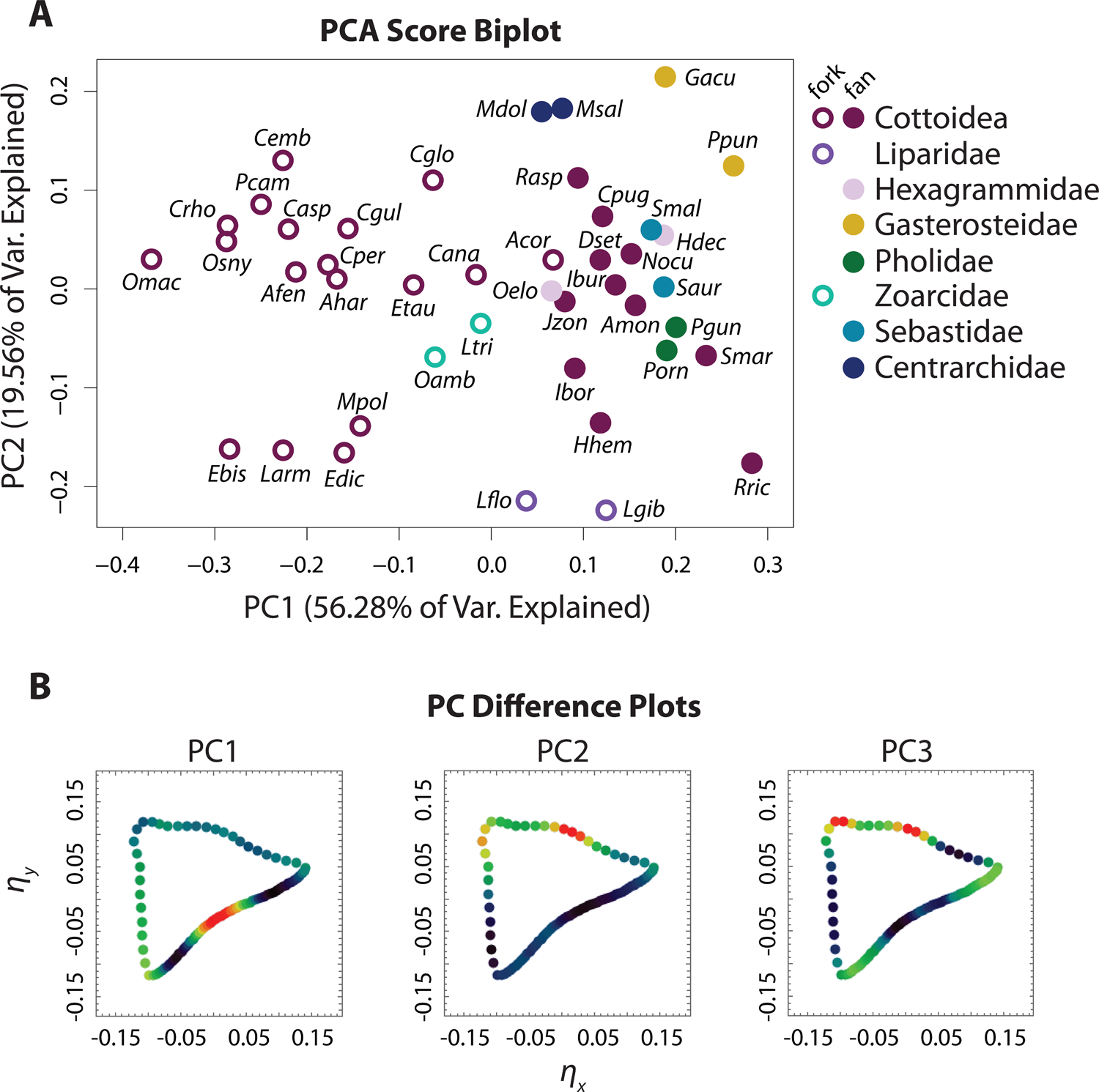
Extended osteoid is a major driver of OP shape variation among cottoid fishes and their relatives. *A*. Eigenshape PCA morphospace (PC1 and 2) of OP shapes only taking mineralized bone into account. Species are coded as either forks (open circles) or fans (closed circles). Note that the major axis of shape variation (PC1) separates fans and forks nearly perfectly. *B*. “Difference plots” for mineralized PCs 1-3, which permit visualization of shape variation “hot spots” in the semi-landmark space (*η*_*y*_ vs. *η*_*x*_) via points that represent semi-landmark coordinate averages. The “hottest” colors (i.e. red) denote outline regions of the greatest shape variation. PC1, accounting for over half of the variance, contributes heavily to diversity in the region that defines the fan-fork continuum. PCs 2 and 3 explain variation independent of the region affected by osteoid, but in regions likely to affect muscle attachment (see Discussion).

PC1 values were strongly associated with the proportion of bone made up of extended osteoid, a quantity calculated from total and mineralized outlines (Fig S5A). This relationship was significant after accounting for phylogeny (Fig S5B; *r*^2^ = 0.8519; df = 38; *p* < 0.001). These insights imply that the presence of extended osteoid is a major feature underlying OP shape distinctions among cottoid species and their outgroups, and that extended osteoid is a primary driver of bone shape likely to influence functional variation of the gill cover.

### Extended osteoid and fork-shaped OPs have evolved multiple times

Based on ancestral state reconstruction using both parsimony and maximum likelihood approaches, a fan-shaped OP is most likely ancestral with respect to the Cottoidei, and a fork-shaped OP has likely arisen multiple times within cottoids (Fig 4A-B). The snailfishes (Family Liparidae) do demonstrate fork-shaped OPs, but there is no evidence for extended osteoid in the center of the bone in this lineage, and they do not fit cleanly into the fan-fork dichotomy. Rather their OPs are “cowboy boot-shaped” (Fig 5), with the notch adjacent to the “boot heel” devoid of extended osteoid. We classified them as forks for the purpose of the ASR but note that the two species we studied (*Liparis gibbus* and *Liparis florae*) occupied a region of PC1 near species with fan-shaped OPs. We also found evidence for derived fork-like OPs outside of the Cottoidei, in two species of eelpout (Family Zoarcidae). The eelpouts we examined (*O. amberensis* and *L. tristichodon*) also demonstrated nonmineralized OP membrane tissue (Fig S2A-B; E), suggesting that osteoid-associated, fork-like OP variants are likely not restricted to cottoid fishes.

**Figure 4.**
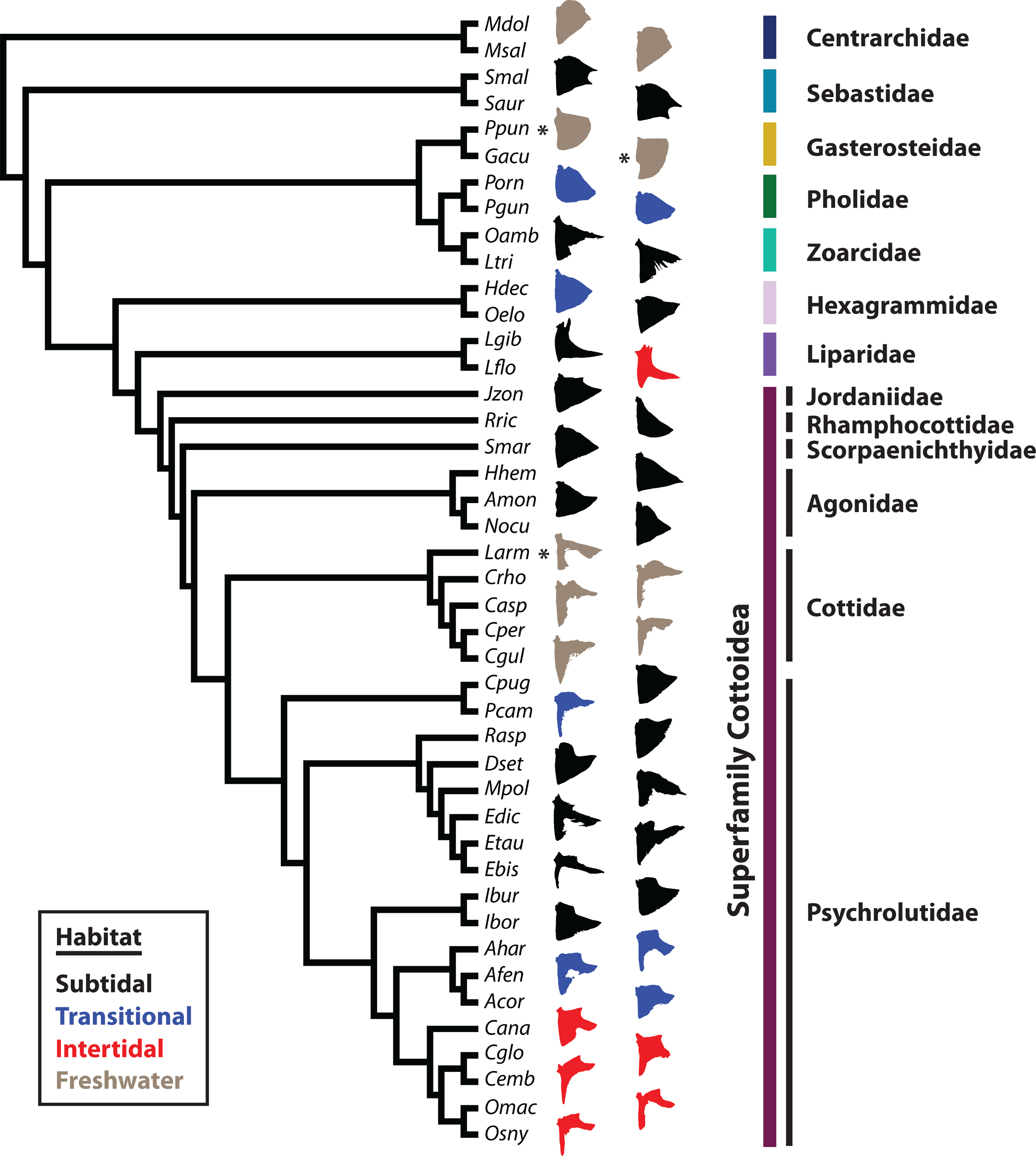
Multiple appearances of extended osteoid during the evolution of cottoid fishes. *A-B*. Ancestral state reconstruction by parsimony and maximum likelihood, respectively, suggesting at least six transitions between fork- and fan-shaped OPs, including the non-cottoid eelpouts (*O. amberensis* and *L tristichodon*). Circles at internal nodes convey the likely character state of the ancestor. In B, marginal likelihoods for either state are represented by color areas within each circle, and in some case suggest that reversals to ancestral fan states were possible. *C*. A continuous analysis of the proportion of the bone area which is extended osteoid. The color scale reflects most likely values for proportion extended osteoid along branches of the tree. Note the similarity in evolutionary pattern between the discrete and continuous metrics of the OP, and that the four “forkiest” OPs (*eelpouts*, *Cottus*, *Porocottus*, and *Oligocottus*) arise from separate lineages.

**Figure 5.**
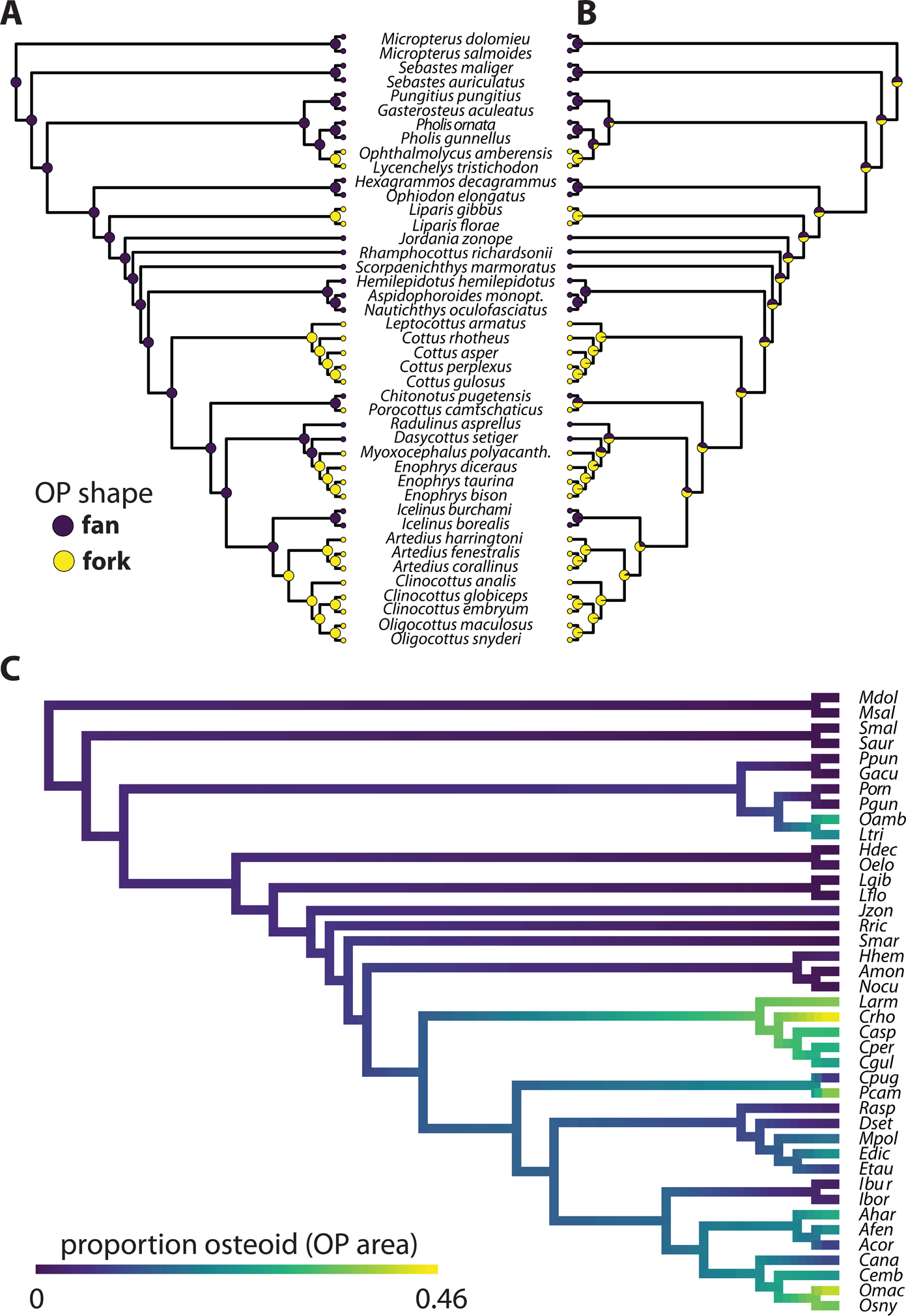
A rich diversity of mineralized OP shape is adorned with fan-fork transitions. Phylogenetic relationships among perciform fish families analyzed in the current study, with a focus on the Cottoidei. Silhouettes are outlines of mineralized (Alizarin-positive) regions from actual bone photographs, although not to scale. Extended osteoid is not shown, but is present in all concave OPs pictured here, except for Sebastidae and Liparidae. Families are colored (vertical bars) according to the scheme from Fig 3. OP silhouette colors correspond to the four main habitats these fish occupy in nature. Species labeled with an asterisk are found in marine or estuarine habitats in addition to freshwater.

Maximum likelihood ASR treating the proportion of bone made up of extended osteoid as a continuous trait also revealed multiple evolutionary events of OP mineralization down-tuning (Fig 4C). Again, we inferred clear cases of large increase in extended osteoid for eelpouts, the lineage leading to sculpin genera *Leptocottus* and *Cottus*, and the tidepool-dwelling sculpin genus *Oligocottus* (Fig 4C; Fig 5). Using phylogenetic generalized least squares and logistic regression we tested whether the proportion of extended osteoid (and also the binary trait of fan/fork) was associated with depth and habitat features, but we found no statistical evidence for an environment-morphology relationship (Table 1; Fig 5).

**Table 1.** OP shape and proportion of extended osteoid are not significantly associated with basic environmental categories reflecting depth and salinity. Shown are results from generalized least squares and logistic regression hypothesis tests in which phylogeny was accounted for. In the two-state analyses environmental factor levels were “deep” and “shallow.” In the three-state analyses environmental factors levels were “deep marine,” “shallow marine,” and “estuary/freshwater.” df = degrees of freedom, and AIC = Akaike Information Criterion. In all cases the null model was a better fit than the model including an effect of habitat.

| **Response variable** | **Model** | **df** | **log likelihood** | **AIC** |
| --- | --- | --- | --- | --- |
| prop. extended osteoid (continuous) | two-state gls full | 3 | 13.5734 | -21.1469 |
|  | two-state gls reduced (null) | 2 | 14.0447 | -24.0894 |
|  | three-state gls full | 4 | 12.7335 | -17.4667 |
|  | three-state gls reduced (null) | 2 | 14.0447 | -24.0894 |
| fan/fork (binary) | two-state logistic full | 3 | -18.8970 | 43.7940 |
|  | two-state logistic reduced (null) | 2 | -19.7567 | 43.5134 |
|  | three-state logistic full | 4 | -18.7270 | 45.4540 |
|  | three-state logistic reduced (null) | 2 | -19.7567 | 43.5134 |

### Extended osteoid appears late in development

How might development have evolved to express extended osteoid and the distinctive fork morphology in the superfamily Cottoidea? One possibility is that bone patterning is reconfigured from the earliest stages of bone ontogeny. On the contrary, our observations of OP form throughout bone development in multiple species revealed that all surveyed OPs began as fans regardless of whether the adult displayed fork or fan shapes, and that large regions of extended osteoid are observed late in bone formation (Fig 6). In species with fan-shaped OPs, fans were present throughout larval and juvenile stages, as illustrated by *Hemilepidotus hemilepidotus* in Fig 6A. Four additional fan species spanning the Cottoidea (*Jordania zonope*, *Rhamphocottus richardsonii*, *Nauthichthys oculofasciatus*, and *Ruscarius meanyi*) also demonstrated this pattern. Conversely, in species with fork-shaped OPs such as *Myoxocephalus polyacanthocephalus*, we did not observe any extended osteoid in the five young larval stages sampled (Fig 6B). The first extended osteoid appeared by 49 days post hatching (d49), and the proportion of osteoid increased progressively (from 6% at d49, to 11% at d80, to 28% in the juvenile for the examples in Fig 6B). The OP of the d224 subadult (last image in the sequence shown in Fig 6B) is a well-defined fork. Fan-shaped OPs also preceded fork-shaped OPs in other fork-bearing species we examined at fewer developmental stages (*Artedius harringtoni*, *Clinocottus acuticeps*, and *Oligocottus maculosus*), representing multiple fan-to-fork transitions within the family Psychrolutidae. Furthermore, in the non-cottoid eelpout species *O. amberensis*, we did not observe a fork-like appearance until later stages of development (Fig S2E). These observations suggest that the fork-fan developmental dichotomy we observed is likely to be general within the superfamily Cottoidea, and very likely beyond, given a similar pattern in the outgroup family Zoarcidae.

**Figure 6.**
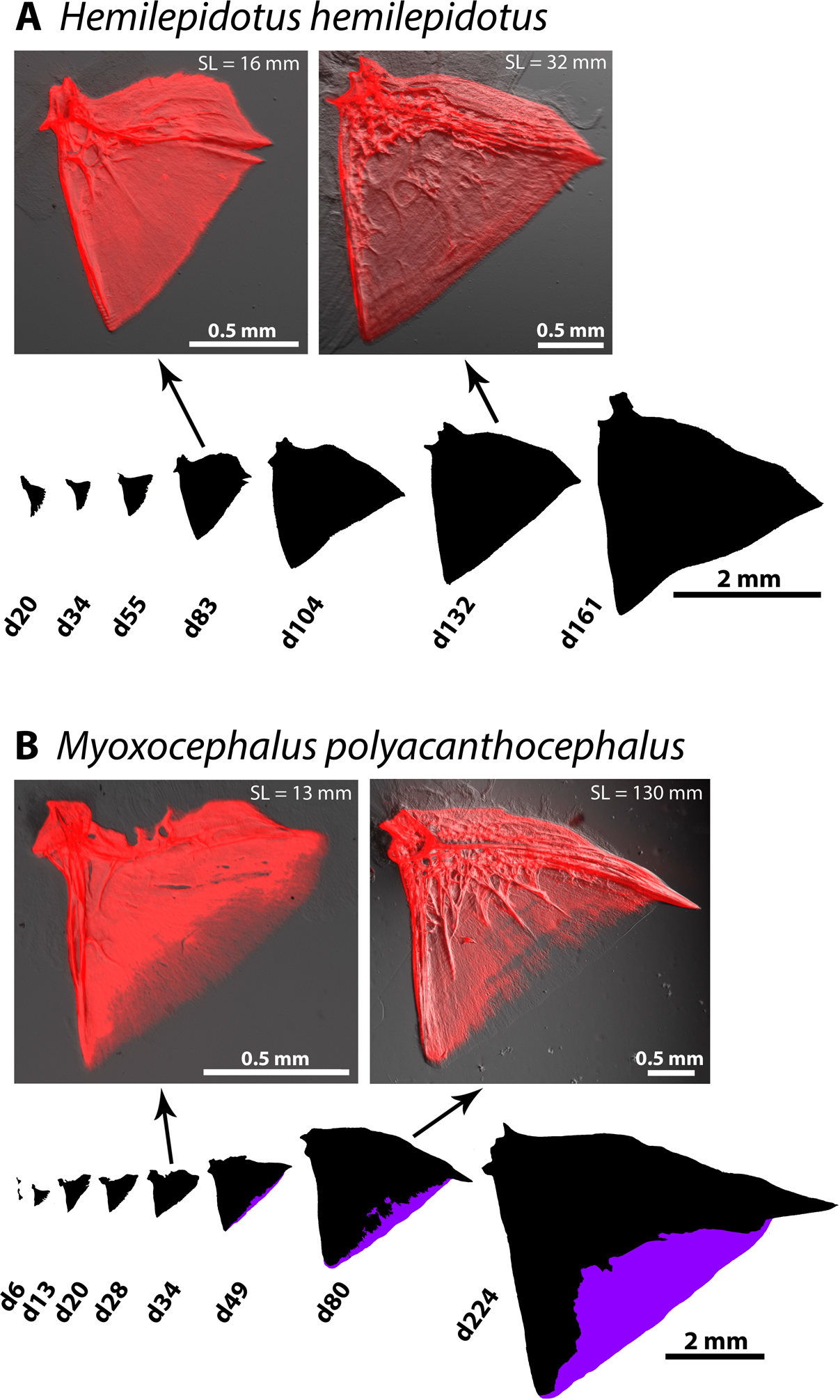
Extended osteoid and the resulting “fork” morphology appear after an established “fan” shape, relatively late in OP development. *A*. A developmental series from a species (*H. hemilepidotus*) with a stereotypical, ancestral “fan” OP, depicted by mineralized bone outlines and Alizarin Red images for two of the outlines. *B*. A similar series for the fork species *M. polyacanthocephalus*. Extended osteoid in *M. polyacanthocephalus* doesn’t appear until 49 days post hatching, and it increases in relative area proportion thereafter.

### The location of extended osteoid changes with bone growth: extended osteoid is not permanent osteoid

Once extended osteoid appears in development is its position stable in the bone as the OP continues growth? We evaluated this question by analyzing overlays of the OPs of different stages, as shown in Fig 7, placing younger OPs on top of older ones. We placed the younger bones near the joint region of older ones (upper left in Fig 7A and 7B), for this is the region where matrix outgrowth begins, as supported by analysis in zebrafish (Kimmel et al. 2010) and the orientation of the incremental banding patterns in sculpins (Fig 2C-D), and other teleosts (Kimmel et al. 2005; Thuong et al. 2017). We observed outgrowth from the joint region toward the OP ventral-posterior edge (lower right), the newest matrix to be formed (see Discussion). For *M. polyacanthocephalus*, the earliest stage shown in Fig 7A is d34, which was before we detected extended osteoid. The first extended osteoid was present at the outgrowing edge at d49, and its location has clearly shifted to the edge of an OP from a later time point (d80), revealing dynamic regulation of osteoid position as the bone grows in a posterior-ventral direction.

**Figure 7.**
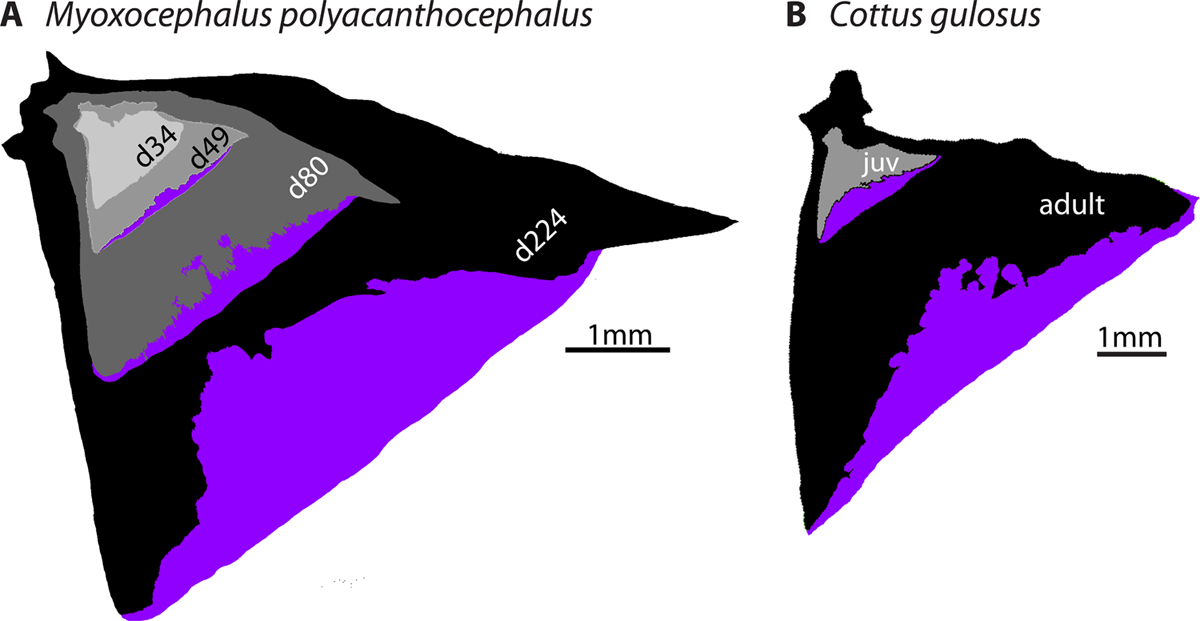
Extended osteoid is not permanent osteoid, but dynamically relocates during OP growth. Multiple stages of OP development for two fork species, superimposed at the joint in the bone outlines to “track” bone positions throughout growth. Clearly extended osteoid during early stages of development eventually becomes mineralized as the bone grows. Extended osteoid patterns in *M. polyacanthocephalus* (*A*.) and *Cottus gulosus* (*B*.).

It was also evident from the overlays that regions of extended osteoid at young stages corresponded to regions of mineralized bone at the later stages. Extended osteoid replacement continued throughout growth; Fig 7B shows such replacement between a young juvenile and an adult of *Cottus gulosus*, an example from the family Cottidae. We note in this example that the OP forms spanning the juvenile-adult transition were quite similar to one another in spite of the marked outgrowth between the two stages, whereas the shape (and proportion of area) of the extended osteoid differed markedly across the larva-juvenile transition sampled in the *M. polyacanthocephalus* series (Fig 7A).

## Discussion

### A novel bone development trajectory widespread among cottoid fishes

We previously surveyed OP variation across 110 teleost families (Kimmel et al. 2017), but a subsequent, expanded focus on the diverse superfamily Cottoidea revealed substantial regions of mineral-deficient, membrane-like tissue to be an unusual feature of OPs in this group. Outside of cryptic graphical references to this tissue type in anatomical line drawings of some sculpin species (Yabe 1985), it had not been previously described. What, we asked, is the material nature of this tissue and its development? Our current study suggests that the flexible membrane filling the posterior ventral portion of fork-shaped OPs is a form of osteoid we refer to as “extended osteoid,” owing to its extended but dynamic presence as nonmineralized matrix. These findings are supported by a variety of analyses, including calcification detection (Alizarin Red staining), material density examination (μ-ct), and histological staining (Ralis-Watkins histochemistry), which all showed that the membrane observed within the OPs of several species in Cottoidea has extremely low radiopacity and calcium content, and is therefore distinct from mineralized bone. The nearly undetectable intensity of Alizarin Red signal in extended osteoid regions was comparable to that in non-bony tissue regions, suggesting extremely low or no calcification. We rule out the possibility that the lack of strong Alizarin Red signal in extended osteoid could be related to the extreme thinness of the tissue, because mineralized regions of larval zebrafish (Kimmel et al. 2010) and sculpin OP regions both show significant Alizarin Red signal well before they are as thick as the unstained extended osteoid regions we observed. This lack of evidence for mineralization, in combination with osteoid-positive Ralis-Watkins histochemistry data, strongly suggest that extended osteoid is not simply thin, mineralized bone.

Data from our cross-polarized imaging and *in situ* hybridization show that the structure and gene expression patterns of extended osteoid warrant its classification as bone matrix. Specifically, the continuity of form between the membrane and mineralized bone within the OP rules out a non-osteogenic tissue. This in-phase growth banding of mineralized bone and extended osteoid tissues (Fig 2D) suggests a common developmental origin, and recent structural studies of developing bone help explain the ordered pattern. A nanometer-level analysis of compact lamellar bone by Reznikov et al. (2018) revealed a defined, hierarchical structure of matrix composed of collagen helices organized into collagen microfibrils, which are further organized into fibril bundles, and ultimately lamellae. According to this model the initial levels of bone matrix structure form independently of mineralization, and would therefore produce an ordered structure for osteoid as well as compact bone. This insight, and demonstrations that circadian rhythms generate periodicity in osteoblast proliferation (Fu et al. 2005; Iimura et al. 2012) and fracture healing (Kunimoto et al. 2016), are consistent with our interpretation of banding patterns as evidence for true osteoid. Furthermore, the clear presence of *sp7*-expressing osteoblasts at the growing edge of the OP extended osteoid (Fig S3) argues for an osteogenic origin. We conclude that extended osteoid develops in conjunction with neighboring osteoid that ultimately follows the more standard mineralization trajectory, and we observe this pattern in multiple cottoid lineages that have evolved OP extended osteoid independently (Fig S4). The developmental tuning that underlies the delayed mineralization of extended osteoid provides a mechanism for the impressive heterogeneity in OP morphology that has evolved in the Cottoidea (Figs 5 and S3).

Osteoid is a well-recognized stage in bone development but is usually temporary (Ducy et al. 1997; Cowles et al. 1998). Indeed, persistent forms of osteoid are known to exist, although generally only pathologically, as in osteomalacia, a multifactorial disease in which osteoid forms normally but fails to mineralize sufficiently (Feng et al. 2013), or due to dietary deficiencies such as in phosphorus-deficient fish (Witten et al. 2016). While lasting, nonmineralized bone matrix has been identified in the fin spines of some blenniid fish species and in the maxilla of the blenniid *Neoclinus blanchardi*, neither of these traits have been thoroughly described, and the thicker, more rigid tissue suggests a different developmental identity (Hastings and Springer 1994). Apart from the aforementioned, the occurrence of non-pathological osteoid within bone structures also containing normally mineralized tissue is, to our knowledge, undescribed.

### Extended osteoid is a major driver of diversity in cottoid gill cover morphology

As extended eigenshape and subsequent principal component analysis demonstrate, fan- and fork-shaped OPs are well segregated within the morphospace along the first principal component axis (Fig 3). Furthermore, the proportion of the OP area made up of extended osteoid co-varied strongly with PC1 (Fig S5). Indeed, the first principal component explains over half of the overall OP shape variation, indicating that variation in extended osteoid among the sampled species forms a major basis for variation in OP shape. This result is significant given the functional importance of the OP as the largest skeletal constituent of the teleost gill cover. The extreme flexibility and thinness of extended osteoid make it morphologically distinct from normally mineralized bone, which suggests that such variation could influence the performance of the gill cover, perhaps in terms of respiration rate, water movement, feeding, gill desiccation prevention, or protection against predators.

### The repeated occurrence of extended osteoid suggests parallel evolution of fork-shaped OPs

Both maximum likelihood and parsimony ancestral state reconstruction suggest 1. that fan-shaped OPs are ancestral with respect to the superfamily Cottoidea, 2. that fork-shaped OPs have arisen multiple times throughout the phylogeny, and 3. that deeply forked OPs have appeared independently at least three times (Fig 5C). One alternative scenario is that one or more “reversals” from the derived fork to the ancestral fan occurred, based on the likelihood analysis. In this case ancestors of 1. *Icelinus*, 2. the clade contaning *Radulinus*, *Dasycottus*, *Myoxocephalus*, and *Enophrys*, and 3. *Chitonotus* would have experienced reversals (Fig 5B). A larger phylogenetic sampling of cottoid species, especially from the scuplin family Psychrolutidae, would be required to more rigorously evaluate this possibility. Other examples of fork-like OPs (Kimmel et al. 2017) exist outside of the species we examined for this study, for example in batfishes (Ogcocephalidae), but without the presence of extended osteoid. Similarly the cottoid family Liparidae (snailfishes) demonstrates fork-shaped OPs with no extended osteoid (see Results). In these cases the developmental processes leading to major OP shape variation are likely very different from those driven by extended osteoid in cottoids and zoarcids. Our evidence suggests that the repeated evolution of fork-shaped OPs in cottoids and zoarcids has occurred via a similar developmental process (extended osteoid), and is therefore an example of “parallel” evolution, at least in the broad sense. Whether the same developmental pathways and individual genes have been modified in parallel among these lineages remains unknown.

Based on our survey, it is clear that a certain evolutionary lability exists for the severity of extended osteoid, and therefore fork-shaped OPs, in the superfamily Cottoidea. Whether this lability is related to a simple and tunable genetic “switch,” repeatedly adjusted during the sculpin radiation by selective means or otherwise, is currently unknown. Future genomic investigations into the genetic basis of OP shape variation are promising and quite possible, as scale (prickle) number in *Cottus* sculpins has been tied to the *ectodysplasin* pathway using QTL mapping (Cheng et al. 2015). Ultimately functional testing, informed by genotype-phenotype association studies in sculpins and/or bone mineralization regulatory networks understood from animal models like zebrafish, will be required to test whether a specific regulatory switch or developmental pathway has been repeatedly modified to tune extended osteoid evolution across the group.

We tested for associations between extended osteoid-driven OP morphological variation and habitat using basic phylogenetic comparative approaches to address the possibility that depth and/or salinity might play an adaptive role in cottoid OP shape evolution. We found no statistical evidence for such associations, but the number of “environmental transitions” across the phylogeny, and our taxon sample size, were both relatively low. Past studies of sculpins have reported significant links between habitat and other traits such as scale number, body size, and body shape (Knope and Scales 2013; Buser et al. 2017). It is therefore possible that an unknown attribute of habitat, life history, or otherwise - one not measured in our study - is fundamentally associated with variation in OP shape, and particularly the multiple fan-fork transitions. It is interesting, for example, that we observed a complete absence of fan-shaped OPs in species living in freshwater or intertidal habitats. Future phylogenetic comparative studies leveraging greater taxonomic sampling and more detailed species-specific information about cottoid biology are warranted.

Regardless of the possible ecological underpinnings, our work suggests that extended osteoid in cottoid fishes is ideal for studying the evolutionary and developmental mechanisms at work in situations of phenotypic convergence. While fan-shaped OPs appear to be ancestral with respect to perciformes and cottoids, the fork shape has emerged repeatedly, both in various sculpin clades, and outside of Cottoidei, in eelpouts. The group provides an interesting model to explore the likely possibility of evolutionary parallelism for a relatively simple trait, a phenomenon observed on a microevolutionary scale in the plate reduction of three-spine stickleback, for example (Cresko et al. 2004; Colosimo et al. 2005), and on a macroevolutionary scale in cases of repeated pigmentation gain or loss (Pointer and Mundy 2008; Rosenblum et al. 2010; Rogers et al. 2013). Cottoid extended osteoid also provides opportunities to identify the gene regulatory pathways involved in the mineralization of osteoid, and to investigate whether these pathways are the same across species and bones, as a similar membrane is present in the IOP, and SOP of many cottoids.

### Dynamic development: replacement of mineralized bony regions with extended osteoid does not occur, but we do observe the opposite

How developmentally malleable is the change in bone matrix between mineralized bone and extended osteoid? Does extended osteoid form in a region of the OP that was mineralized at an earlier stage? The flip side of this question is to ask whether a region of extended osteoid can later become mineralized. Both questions address the basic issue of developmental plasticity (and heterogeneity in form) of the bone matrix in the fork species. We looked at the distribution of extended osteoid over developmental time in multiple fork-bearing species, assisted by superimposition of bones from different developmental stages (Fig 7). We found that once a region is mineralized, it does not return to a state of extended osteoid, eliminating a “de-mineralization” mechanism from explanation. However, as the OP grows outward, regions previously occupied by extended osteoid are eventually mineralized. Successive addition of osteoid then continues at the new edge, past that of the previous stage. Either one form of the matrix or the other is laid down at the growing OP edge, depending on the species of fish.

We interpret this pattern to mean that extended osteoid, although present for a much longer period of time than osteoid in fan-bearing species, is not permanently maintained during development. Maintenance of the OP shapes during juvenile development would of course be impossible without exquisite and dynamic regulation of extended osteoid – added at the growing bone edge, and at the same time overwritten by mineralization in the older matrix. Higher resolution developmental and genetic studies in the future will elucidate this dynamic developmental process, and to what extent it may vary among different lineages.

### Multilevel heterochrony: forks represent a derived, late-appearing instance of cellular paedomorphosis

We observed that in both fan- and fork-bearing species, the OP develops initially as a fan, but in fork-bearing species extended osteoid is then expressed at a later stage, presumably by blocking mineralization or by undergoing delayed mineralization. Subsequently, with continued bone growth the fan morphology changes to a fork. This temporal pattern of extended osteoid development supports a classical argument made by von Baer in the early 19^th^ Century (von Baer 1828), and then extended by Darwin (Darwin 1859), that an evolutionarily ancestral state precedes an evolutionarily derived (specialized) state in development. Based on our ASR we conclude that this pattern is consistent with cottoid OPs: the early developmental fan state is ancestral, and the following fork state possessing extended osteoid is derived.

Furthermore, considering that presence of osteoid typically marks a transient and brief early stage of bone development in vertebrates, we interpret this new pattern of delaying and lengthening the period when osteoid is prominent as indicative of heterochrony – a change in developmental timing between ancestor and descendant (Alberch et al. 1979; Klingenberg 2007). Our case seems to fit only poorly or not at all into the classical scheme of heterochrony, where changes are considered either as paedomorphic or peramorphic: In paedomorphosis, later developmental stages in the descendant retain characteristics of ancestral early stages, clearly not the case here for fork-species because the ancestral early stages are fans, not forks. In peramorphosis, development in descendants is pushed beyond the end of an ancestral developmental sequence, producing a somehow exaggerated ancestral form such as the famous case of the Irish elk, possessing antlers much larger than its ancestor (Gould 1977). These definitions are, however, ill-suited to extended osteoid and fork-like OPs, because they refer to development strictly at the *organismal* level. Delayed mineralization, in the case of extended osteoid, occurs at the *cellular* level. Here, the extended osteoid cells (osteoblasts) retain characteristics of early (pre-mineralization) states, but this phenomenon is nested within, and indeed occurs as a later feature of, organismal development. In other words, the derived phenotype (fork-shaped OPs) is the result of cell-level paedomorphosis that begins at later developmental stages in fork-bearing species, but never initiates in fan-bearing species. A fish with fork-shaped OPs begins its development with osteoblasts that achieve mineralization quickly, but by the time the individual is finished developing into an adult, many of the osteoblasts composing its OPs experience a delay in their own cellular development. In this sense, the multilevel heterochronic changes in cottoid OPs have produced a form that is not at all like ancestral early or late stages, but novel. A systems-level understanding of this particular form of novelty will ultimately require precise estimates of heterochronic shift variation among lineages with fork-shaped OPs, in addition to molecular interrogation of the cell-level gene regulatory processes that ultimately underlie extended osteoid.

## Summary

We describe a new developmental trajectory for craniofacial bones in fishes based on histological, tomorgraphic, and molecular evidence. The resulting tissue state, which we call “extended osteoid,” and the developmental down-tuning of mineralization that presumably gives rise to it appear to be a heretofore undescribed phenomenon in evolutionary biology, and extended osteoid helps to explain a large proportion of the morphological diversity among opercle bones of bony fishes in the suborder Cottoidei. This membranous tissue has evolved several times in (and at least once outside of) cottoid fishes, although no ecological bases for the pattern of convergence are yet clear. The repeated appearance of similar morphologies in distinct species offers evolutionary developmental biologists a unique opportunity to test which aspects of the genotype-phenotype map are repeatable on a macroevolutionary scale. Extended osteoid in the cottoids also holds promising potential as a model for studying osteogenesis, given biological separation of the processes of bone formation and its mineralization in a non-pathological context. We propose that variation in extended osteoid within and among various species of sculpin offers a potentially useful window into the genetics of mineralization-related bone pathologies such as osteomalacia, and of repeated evolution of form via developmental processes in general.

## Acknowledgments

We thank Kevin Clifford (Oregon Coast Aquarium) and members of the Kimmel and Cresko Labs at the University of Oregon for specimens, assistance with specimen collection, and/or helpful advice on the direction of this study. TD thanks the captain and crew of the ARSV Laurence M. Gould and personnel of the U. S. Antarctic Program for assistance with eelpout collections. N. MacLeod made available Mathematica workbooks for the extended eigenshape analysis. This work was supported by grants from the National Science Foundation (IOS-0818738 to CBK and PLR-1543383 to TD), the National Institutes of Health NIDCR (R01 DE13834 to CBK), and the NIH NICHD (P01-HD022486 to J. Eisen). CMS was supported by NIH NCRR grant RR032670 to W. Cresko. The authors declare no conflicts of interest.

## Author Contributions

EC, CMS, and CBK conceived and designed the study, all authors participated in data collection, and CMS and CBK performed the statistical analyses. EC, CMS and CBK drafted early versions of the manuscript and all authors contributed to later versions of the manuscript.

## Data Accessibility

Upon publication all summary data (PC values, proportion osteoid estimates, etc.) and bone images will be available in the Dryad Digital Repository.

